# THERAPEUTIC EFFECTS OF HYPOXIC AND PRO-INFLAMMATORY PRIMING OF MESENCHYMAL STEM CELL-DERIVED EXTRACELLULAR VESICLES IN INFLAMMATORY ARTHRITIS

**DOI:** 10.1101/2021.06.28.450178

**Authors:** Alasdair G. Kay, Kane Treadwell, Paul Roach, Rebecca Morgan, Rhys Lodge, Mairead Hyland, Anna M. Piccinini, Nicholas Forsyth, Oksana Kehoe

**Affiliations:** Department of Biology, University of York, York, UK; School of Medicine, Keele University at the RJAH Orthopaedic Hospital, Oswestry, UK; Chemistry Department, Loughborough University, Loughborough, UK; School of Chemistry, University of Nottingham, Nottingham, UK; School of Pharmacy, University of Nottingham, Nottingham, UK; School of Pharmacy and Bioengineering, Keele University, The Guy Hilton Research Laboratories, Hartshill, Stoke on Trent, UK

**Keywords:** Rheumatoid Arthritis, inflammation, immunomodulation, extracellular vesicles, mesenchymal stem cells

## Abstract

Novel biological therapies have revolutionised the management of Rheumatoid Arthritis (RA) but no cure currently exists. Mesenchymal stem cells (MSCs) immunomodulate inflammatory responses through paracrine signalling, including via secretion of extracellular vesicles (EVs) in the cell secretome. We evaluated the therapeutic potential of MSCs-derived small EVs in an antigen-induced model of arthritis (AIA).

EVs isolated from MSCs cultured normoxically (21% O_2_, 5% CO_2_), hypoxically (2% O_2_, 5% CO_2_) or with a pro-inflammatory cytokine cocktail were applied into the AIA model. Disease pathology was assessed post-arthritis induction through swelling and histopathological analysis of synovial joint structure. Activated CD4+ T cells from healthy mice were cultured with EVs or MSCs to assess deactivation capabilities prior to application of standard EVs *in vivo* to assess T cell polarisation within the immune response to AIA.

All EVs treatments reduced knee-joint swelling whilst only normoxic and pro-inflammatory primed EVs improved histopathological outcomes. In vitro culture with EVs did not achieve T cell deactivation. Polarisation towards CD4+ helper cells expressing IL17a (Th17) was reduced when normoxic and hypoxic EV treatments were applied in vitro. Normoxic EVs applied into the AIA model reduced Th17 polarisation and improved Th17:Treg homeostatic balance.

Priming of MSCs in EV production can be applied to alter the therapeutic efficacy however normoxic EVs present the optimal strategy for broad therapeutic benefit. The varied outcomes observed in MSCs priming may promote EVs optimised for therapies targeted for specific therapeutic priorities. EVs present an effective novel technology with potential for cell-free therapeutic translation.

## Introduction

Mesenchymal stem cells (MSCs) are a promising therapeutic option owing potential for tissue repair through trilineage differentiation capacity, immunomodulatory properties disrupting T cell proliferation, B cell function and dendritic cell (DCs) maturation; and promoting anti-inflammatory responses mediated through macrophage interactions^1^. Widespread introduction of stem cell therapies has been hindered by inconsistent outcomes at clinical trial and donor variability. Our group has demonstrated the immunomodulatory capacity of both MSCs and their conditioned medium (CM-MSC) to reduce inflammation in a murine antigen-induced arthritis (AIA) model through enhanced Treg function and restored Treg:Th17 ratio^2,3^. MSCs convey their immunomodulatory properties through cell-to-cell contact, autocrine responses and paracrine signalling^1^, including through secretion of extracellular vesicles (EVs)^4^. EVs are membrane bound particles that carry a cargo of microRNA (miRNA), mRNA, lipid, carbohydrate and protein signals to facilitate intercellular communication^5–9^. Notably, disease severity in seropositive RA has been linked to EVs signalling^10–15^. MSC-derived EVs have been shown to be beneficial in autoimmune disorders in functioning to modulate autoimmune responses, particularly related to graft rejection and hypertension^16–18^ but also in inflammatory arthritis and rheumatic diseases^19–21^. T cells have been shown to recruit EVs released by DCs, suggesting an integral action in orchestrating *in vivo* immune responses^22,23^. Furthermore, tolerance in DCs is influenced by CD4+ T cells through EVs-mediated intercellular signalling mediated by transfer of EVs-packaged miRNA^24^. We hypothesised that EVs recapitulate the anti-inflammatory and immunomodulatory effects of MSCs in the arthritis model and that priming MSCs during the generation of EVs through hypoxic culture or pro-inflammatory cytokine cocktails will enhance therapeutic outcomes.

Identification of the most effective cell source and maintenance conditions for generation of optimally efficacious EVs will accelerate development of EV therapeutics for RA and other regenerative medicine targets. To date, no research has directly examined and contrasted the influence of cell isolation and priming on EVs generation using both hypoxia and pro-inflammatory pre-conditioning of MSCs during production.

Here we present novel data utilising priming strategies for MSC-derived EVs applied to the AIA model of inflammatory arthritis. We investigate amelioration of symptoms through reduced swelling and histopathological improvement, and EVs influence on T cell proliferation and polarisation *in vitro* and *in vivo.* We hypothesise that EVs represent a potential therapeutic approach for the treatment of inflammatory arthritis that may encounter less obstacles than cell therapy to widespread application in the clinic.

## Results

### EVs isolated through differential ultracentrifugation characterise as EVs according to internationally agreed criteria

All EVs isolated in this study were enriched using 100,000 x g ultracentrifugation with EV pellets resuspended based on counts of source cells taken at the point of collection, with resuspension of vesicles from 1.0 × 10^6^ MSCs per 30μl.

A combination of quantitative and qualitative analyses was performed to confirm isolation of EVs. EV preparations from MSCs cultured under normoxic conditions were assessed by flow cytometry analysis using the MACSPlex exosome kit (Miltenyi Biotec) to identify 37 exosomal surface epitopes, including characteristic exosome markers CD9, CD63 and CD81. EV isolations showed elevated enrichment in mean expression values for CD9 (81.25±5.03%), CD63 (94.59±2.23%) and CD81 (79.41±9.07%) (n=11) (representative example Figure S1A) confirming the identity as EVs. Western blot analysis of MSC lysate and EVs (EV-NormO2) determined positive detection of Alix, a transferrin receptor binding protein involved in multivesicular body (MVB) biogenesis. MVBs are a late endosomal compartment functioning to traffic biomolecules in the form of intralumenal vesicles to either lysosomes for degradation or to the plasma membrane for extracellular secretion^25^. Western blot also confirmed the absence of cytochrome C, an ubiquitous mitochondrial protein acting as negative control (Figure S1B)^26^. Transmission Electron Microscopy (TEM) imaging demonstrated the presence of spherical vesicles in isolated preparations of normoxic and hypoxic EVs in accordance with international standards for single EVs characterisation (Figure S1C). Finally, EVs were characterised for their size distribution and concentration using the Nanopore technology (Izon Science). EV preparations showed a distribution of EV sizes with most prevalent diameter of ~ 200 nm with maximal diameter ~500 nm (Figure S1D-E). Together, these results demonstrate that our differential ultracentrifugation methodology successfully isolates EVs according to internationally agreed criteria. We observed that pro-inflammatory priming of MSCs using the methodology adopted here was possible for a period of 48 hours for collection of CM-MSC, but extended cell culture in the presence of pro-inflammatory cocktail to 72 or 96 hours resulted in consistent reductions in cell numbers below the originally seeded cell numbers, suggestive of induced crisis in the cultured cells (data not shown).

### Priming of MSCs does not affect total protein content in EV cargo

We set out to determine whether the total protein content in the cargo of EVs was impacted by cell priming. A 4 parameter polynomial nonlinear regression of log-transformed bicinchoninic acid assay (BCA, Pierce Biotechnology) data revealed no significant increase in total protein between conditions EV-2%O2 preparations (81.29±34.82 pg/1.0 × 10^6^ cells; n=8) and EV-Pro-Inflam (69.09±37.38 pg/1.0 × 10^6^ cells; n=9) compared to EV-NormO2 (40.43±14.73 pg/1.0 × 10^6^ cells; n=11) (p>0.05).

### Application of MSC-derived EVs ameliorates histopathology and joint swelling in AIA

To assess the therapeutic potential of MSC-derived EVs we introduced each of our primed MSCs EVs treatments into a murine model of inflammatory arthritis and measured effects of treatments on histopathological outcomes. AIA is an acute model of inflammatory arthritis that typically exhibits peak joint swelling at 24 hours post induction with clinical symptoms and histopathological signs that resemble rheumatoid arthritis. We have previously demonstrated the efficacy of MSCs and CM-MSC in ameliorating swelling in AIA^2,3^. In this study, we compared three EV treatments to PBS only vehicle controls, measuring the reduction of joint diameter from peak swelling (day 1). Local administration of EVs into joints significantly reduced joint diameter post-arthritis induction, a quantitative measure proportional to joint swelling. Specifically, EV-NormO2 (day 2 = 4.4±0.6mm, day 3 = 8.0±0.6mm, p<0.01), EV-2%O2 (day 2 = 7.7±0.8mm, day 3 = 11.0±0.9mm, p<0.001) and EV-Pro-Inflam (day 2 = 6.7±1.0mm, day 3 = 11.3±0.9mm, p<0.001) all very significantly reduce joint swelling in comparison to PBS vehicle control treatments (day 2 = 0.2±0.8mm, day 3 = 2.8±1.0mm) (Figure 1A). Reductions in joint diameter from peak swelling were proportional for all treatment conditions.

**Figure 1.**
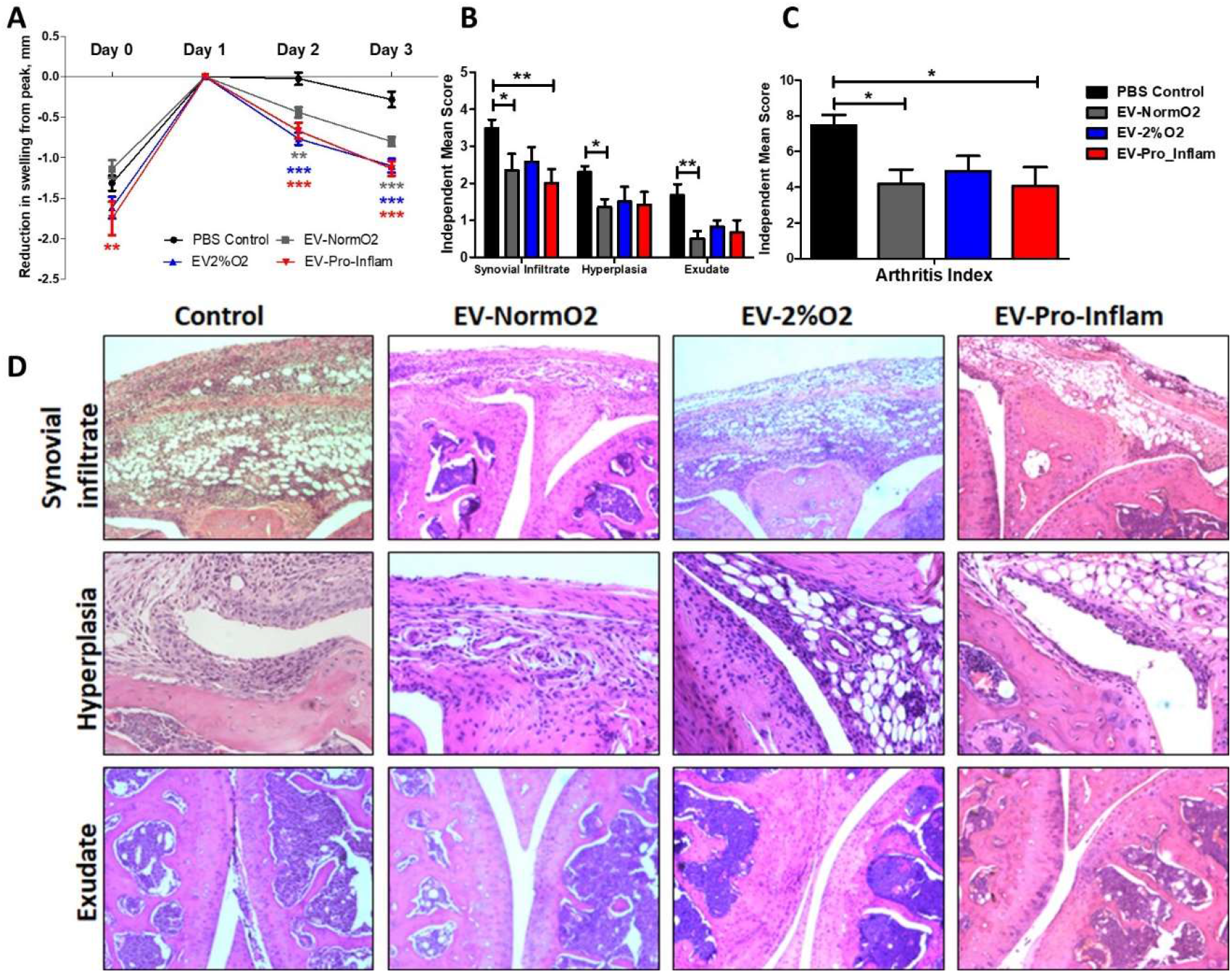
MSC-derived EVs treatment of mice with AIA. (A) Alleviation of joint swelling as a measure of therapeutic efficacy following EV treatments shows a significant effect of EVs compared to vehicle controls at day 2 and day 3 after arthritis induction; normalised to 0 at peak swelling (day 1) (2 Way ANOVA with Repeated Measures and Bonferroni multiple comparisons test post hoc) (B) Examination of histological signs of arthritis pathogenesis following EV treatment shows significant therapeutic effects of EVs sourced from MSCs cultured under normoxia and MSCs cultured in the presence of pro-inflammatory cytokines 3 days post-induction. (C) Combined score arthritis index shows significant reductions in arthritis index 3 days after induction when treated with EVs (1 way ANOVA with Repeated Measures and Bonferroni Multiple Comparisons test post hoc); (D) Representative images for synovial infiltrate, hyperplasia of the synovial lining and synovial exudate into joint cavity for vehicle control and EVs treatments (Control n=21, EV-NormO2 n=8, EV-2%O2 n=6, EV-Pro-Inflam n=6. *p<0.05; **p<0.01; ***p<0.001)

Histopathological symptoms include immune cell infiltration into the synovium; hyperplasia of the synovial membrane; and extravasation of leukocytes into the synovial joint cavity. We have previously shown that the intra-articular injection of MSC conditioned medium in mice with AIA ameliorates damage as observed through histopathological analysis^3^. Whilst EV-2%O2 overall showed a tendency to reduce total arthritis index (AI) scores (4.92±0.84) compared to PBS controls, this was not statistically significant. For EV-NormO2, this reduced AI score reflected decreases from control scores in hyperplasia of the synovial membrane (1.35±0.22 vs 2.30±0.17) and joint exudate (0.50±0.21 vs 1.68±0.30), whilst EV-Pro-Inflam demonstrated reduced synovial infiltrate (2.0±0.39 vs 3.49±0.24) (Figure 1B). Overall AI showed significant reductions in mice treated with EV-NormO2 (4.20±0.79) and EV-Pro-Inflam (4.08±1.04) compared to PBS control (7.46±0.59) (Figure 1C). Taken together, these results demonstrate EV-NormO2 treatment produced a broad reduction in swelling and improvement in histological outcomes, whilst primed MSCs were effective in reducing swelling, but evidence suggested this was not exclusively through reductions in synovial infiltrate or reduced synovial damage.

EV-Pro-Inflam treatments showed reductions in synovial infiltrate sufficient to affect a significantly improved overall arthritis index, however this treatment methodology was not as effective at reducing synovial exudate or hyperplasia of the synovial membrane (Figure 1D). The reduction in synovial infiltrate may therefore contribute towards reduced swelling however the results seen in EV-2%O2 suggests this is not a single factor underpinning the mechanism of action.

### EV treatments do not affect expression of pro- (TNF-α) and anti-inflammatory (IL-10) cytokines detectable in serum of AIA mice

The pro-inflammatory cytokine TNF-α is a key driver of disease pathogenesis in RA and a therapeutic target in biological treatments^27,28^. Conversely, IL-10 is a master regulator of anti-inflammatory immune responses^6^. To investigate the effects on circulating cytokines, TNF-α and IL-10 were measured at day 3 in the serum of mice following EV treatments. The circulating concentration of TNF-α in serum of treated mice did not vary significantly between controls (7.02±0.47pg/mL, n=15) and EV treated conditions or between treatments EV-NormO2 (7.38±1.04pg/mL), EV-2%O2 (6.93±0.65pg/mL) or EV-Pro-Inflam (5.73±0.47pg/mL) (p>0.05, n=5 per treatment). ELISA on mouse serum could not detect IL-10 in treatment or control conditions (data not shown). Owing to the small volume of synovial fluid in the murine articular cavity, it was not possible to isolate sufficient synovial fluid to accurately detect cytokine expression within the localised joint cavity. The consistent concentrations of circulating TNF-α in serum of mice undergoing treatment and in untreated control mice suggests that immunomodulation of TNF-α regulated cytokine/chemokine induction is not a primary mechanism underpinning efficacy of EVs treatments in reducing swelling and/or histological improvement. This is in contrast to results previously observed in MSC treatments^2^.

### EVs with hypoxic priming or unprimed decrease Th17 polarization but only MSCs suppress proliferation when co-cultured with activated CD4+ T cells in vitro

Prior to introduction *in vivo*, we investigated the *in vitro* effect of MSC-derived EVs on T cell polarisation and proliferation to establish an alternative mechanistic rationale for efficacies observed in swelling and histological scores for EVs treatments. MSCs have been shown to deactivate T cells in co-culture^3^. MSCs and EVs produced following conditions EV-NormO2, EV-2%O2 and EV-Pro-Inflam were therefore co-cultured for 5 days with activated T cells isolated from healthy mice and polarisation and proliferation of activated cells assessed as a measure of immune response and T cell deactivation.

T cell polarisation was assessed by flow cytometry analysis of surface CD4 and intracellular markers characteristic of Th1 (IFN-γ), Th2 (IL-4) and Th17 (IL-17a). The ability of EV treatments to affect deactivation of T cells, and therefore influence T cell proliferation and extent of immune response, was examined to determine experimental conditions for assessment of EV treatment interactions with T cells *in vivo.*

Replicating previous studies^3^, MSC with T cell co-cultures prompted significantly increased numbers of CD4+ T cells compared to activated T cells cultured alone. MSCs also elicited increased CD4+ T cells in comparison to EV-2%O2 and EV-Pro-Inflamm treatment conditions (n=3, p<0.05) but not EV-NormO2 (Figure 2A).

Further analysis of IL17a expressing cells within the CD4+ population demonstrated that MSC co-cultures increased cell proportions in comparison to PBS controls and all EV conditions, whereas all EV treatments showed a decrease in Th17 cell polarisation compared to MSCs treatment (p<0.05, n=3), with both EV-NormO2 and EV-2%O2 treatments showing significantly reduced IL-17a-expressing cells compared to PBS controls (p<0.001, p<0.05 respectively) (Figure 2B). Whilst no differences were seen for IFN-γ expressing cells (Th1) (Figure 2C), IL-4 expressing cells (Th2) were elevated in MSC co-cultures (6.00±0.24%) compared to EV-Pro-Inflam (2.74±0.52%) (p<0.01); EV-NormO2 (2.86±0.30%) or EV-2%O2 (3.22±0.03%) but not in comparison PBS control (4.79±0.99%) (p>0.05, n=3). Notably, EV-NormO2 and EV-Pro-Inflamm treatments reduced proportions of IL4-expressing T cells (Th2) in comparison to PBS controls (p<0.05) (Figure 2D). Consequently, pro-inflammatory cocktail priming of MSCs generated EVs that reduced “anti-inflammatory” IL-4-expressing T cells (Th2) but did not reduce IL-17a-expressing, pro-inflammatory T cells (Th17). Statistical analysis was performed using log transformed data.

**Figure 2.**
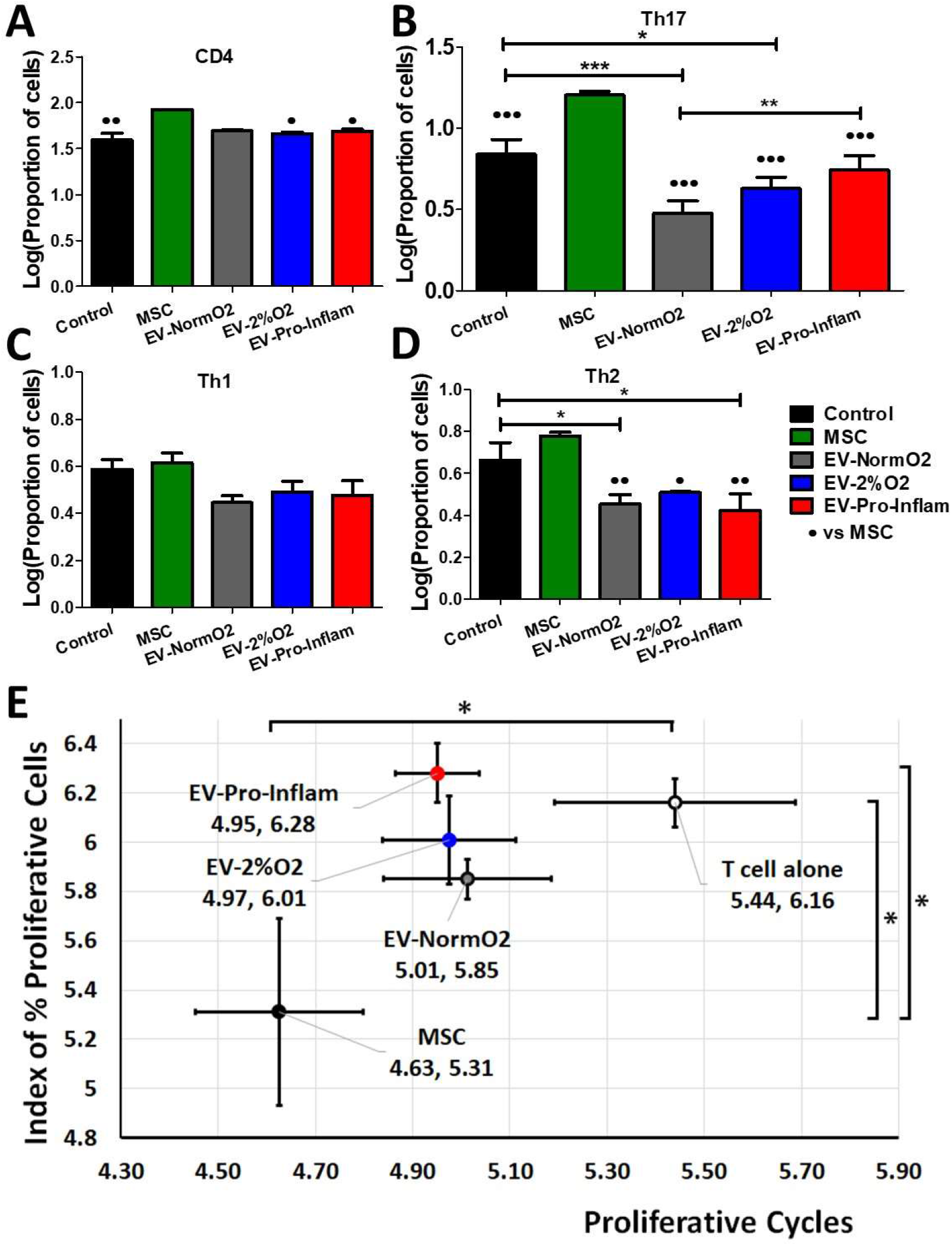
Outcomes of EV treatments co-cultured with T cells isolated from healthy murine spleens. **(A)** Increased CD4+ T cells in MSC co-cultures compared to EV-2%O2; EV-Pro-Inflam; and T cells alone control **(B)** Increased pro-inflammatory Th17 cells (IL17a+) in MSC co-cultures over to EV-NormO2; EV-2%O2; EV-Pro-Inflam; and T cells alone control with EV-NormO2 significantly reduced on PBS control and EV-Pro-Inflamm also **(C)** EV or MSC treatments did not alter Th1 polarisation **(D)** EV treatments all reduced Th2 polarisation in comparison to MSC treatment, with EV-NormO2 and EV-Pro-Inflam also reduced in comparison to PBS controls (Index of proliferation only) (n=3; *p<0.05; **p<0.01; ***p<0.001; 1-Way ANOVA with Bonferroni post-hoc of log transformed data. **(E)** MSC co-cultures significantly inhibited T cell proliferation compared with T cells cultured alone (both measures) and EV-Pro-Inflam co-culture (Index of proliferation only) (n=3; *p<0.05; **p<0.01; ***p<0.001; 1-Way ANOVA with Bonferroni post-hoc of log transformed data.

Additionally, the expression of IL17a per cell, represented by mean fluorescence intensity (MFI) of signal, was significantly reduced in EV-NormO2 (3311.74±33.71) and EV-2%O2 T cell co-cultures (3367.19±39.92) compared to CD4+ T cells cultured alone (3728.69±50.28) (p<0.01, p<0.05, n=3). The impact of this change would need to be assessed *in vivo* as this had not translated to reduced Th17 numbers *in vitro.*

These results demonstrate that *in vitro* MSC-derived EVs do not prompt increases in CD4+ T cell numbers or Th17 effector cell polarisation as seen with MSC treatments. Interpretation of this data is limited by the lack of Treg data preventing the examination of the Treg:Th17 ratio which is offset in RA.

We have previously shown the ability of MSCs to suppress T cell proliferation in co-culture, with MSCs being more effective than CM-MSC^3^. MSC with T cell co-culture reduced the proportion of proliferating T cells, measured via a proliferative index^29^ and the number of proliferative cycles^30^ undergone by T cells in co-culture (5.31±0.38, 4.63±0.17 respectively) in comparison to T cells cultured alone (6.16±0.10, 5.44±0.25) (n=8, p<0.05). In contrast, CD4+ T cells from healthy (not AIA induced) mice showed no significant inhibition of proliferative cycles when cultured with EV-NormO2 (5.85±0.08, 5.01±0.17), EV-2%O2 (6.01±0.18, 4.97±0.14) or EV-Pro-Inflam (6.28±0.12, 4.95±0.09) (n=8, p>0.05) (Figure 2E).

EV treatments showed greater impact on proliferative cycles than on the proliferative index, though treatments did not significantly vary from either MSC co-cultures or T cells cultured alone, with the exception of pro-inflammatory primed EVs which allowed a higher proliferative index in comparison to MSC co-culture (n=8, p<0.05) (Figure 2E). Previous studies have shown similar results through the application of whole secretome^3^. The results found here suggest that direct deactivation of T cells and immunomodulated T helper polarisation does not provide a primary mechanism for therapeutic action of EVs *in vivo.* Whilst MSCs do display direct immunomodulation of T cells, it may be that MSC-secreted EVs require an intermediate such as DCs to affect similar influence on T cell responses.

### MSC-derived EVs reduce IL-17a-expressing T cell polarisation and restore the Th17:Treg balance ex vivo

Following on from our *in vitro* study of T cell deactivation using CD4+ T cells of healthy mice, we hypothesised that direct T cell interactions were not likely to be the mechanism through which therapeutic effects are observed following EVs infusion. EV-Pro-Inflam treatments did not influence T cell polarisation however hypoxic priming of MSCs generated EVs capable of influencing Th2 polarisation and Th17 expression (observed through increased mean fluorescence intensity). At this stage, data suggested MSC priming (either 2%O2 or pro-inflammatory cocktail) conveys no beneficial advantage over normoxically produced MSC-EVs in modulating T cell responses or in reducing swelling and affecting improved histological outcomes. Our previous study demonstrated the ability of whole secretome conditioned medium from normoxically grown MSCs to influence T cell polarisation, restoring Treg:Th17 homeostasis, and to affect reductions in swelling and tissue damage *in vivo*, similarly not via T cell deactivation when investigated *in vitro*^3^. We apply the AIA model to examine T cell interactions in only EV-NormO2 vesicles comparative to PBS treated controls. Here we aim to elucidate whether EVs affect T helper cells deactivation *in vivo.* Spleens and lymph nodes (inguinal and popliteal) of EV-NormO2 treated AIA mice and PBS controls were dissociated, and CD4+ T cells isolated. These cells represent *in vivo* primed T cells within the inflammatory arthritis environment prior to EVs treatment, isolation and subsequent *in vitro* activation.

EV-NormO2 did not increase the proportion of CD4+ T cells in spleen (13.28±0.51%) or lymph nodes (15.04±1.04%) over PBS controls in spleen (14.21±1.07%) or lymph nodes (15.52±1.15%) (p>0.05, n=4) (Figure S2A, S2B shown with log transformed data).

Examination of *in vivo* primed CD4+ T cells activated and cultured *in vitro* for 4 hours in the presence of a membrane transport blocker facilitated evaluation of T cell polarisation. These results demonstrated MSC polarisation of T cells favoured IL-17a expressing T cells (Th17) outcomes as shown previously^3^. RA (and other autoimmune disease) sufferers experience an imbalanced Th17:Treg ratio leading to inappropriate immune responses and tissue damage^31–34^ so for the *in vivo* analysis we additionally examined regulatory T cells (Tregs, CD4+CD25+FOXP3+) polarisation to allow calculation of ratios of Th17:Treg to highlight differential changes in polarisation of subsets of helper T cell that could affect the overall homeostatic balance. Percentage data was log transformed for analysis with unpaired T tests (figures show log transformed data).

When compared to PBS control (3.23±0.81%), spleen T cell polarisation towards IL17a-expressing T (Th17) effector cells following EV-NormO2 (0.83 ± 0.10%) treatment showed a significant decrease in the proportion of pro-inflammatory Th17 cells induced in AIA mice (p<0.01, n=4), with an additional significant decrease in CD4+CD25+FOXP3+ (Treg) proportions in EV-NormO2 treated T cells (0.97 ± 0.05) compared to PBS controls (1.12 ± 0.02) (p<0.05, n=4) (Figure 3A). This translated to significantly improved Treg:Th17 ratios in spleens of EV-NormO2 (12.06±2.12:1) treated mice compared to PBS controls (5.03±1.40:1) (p<0.05, n=4) (Figure 3B). This result indicates that EVs can prompt an advantageous shift in the Treg:Th17 balance towards a healthy state (EV-NormO2 vs PBS control, 12.06 ± 2.12 vs 5.03 ± 1.40, p<0.05, n=4), demonstrating a similar efficacy of applying EVs alone to that previously seen when applying CM-MSC^3^. A similar trend was seen in T cells from lymph nodes with a significant reduction in Th17 polarisation in EV (0.90±0.09%) compared with PBS controls (4.26±1.12%) (p<0.05, n=4) (Figure 3C) leading to improved Treg:Th17 ratio (EV-NormO2 vs PBS control, 9.01 ± 0.80 vs 3.78 ± 1.80, p<0.05, n=4) (Figure 3D). Taken together, these data show that EV treatments were capable of affecting alleviation of symptoms of inflammatory arthritis through reduced Th17 polarisation and shift in the Treg:Th17 balance. We also observed an increase in IFN-γ expressing (pro-inflammatory Th1) T cells in spleens of EV-NormO2 treated mice (6.38±0.40%) compared to PBS controls (4.00±0.35%) (p<0.05, n=4) (Figure S2C). Although no change was observed in IL-4 expressing (Th2) T cells (Figure S2D), increased Th1 polarisation lead to an increased Th1:Th2 balance in EV-NormO2 treated mice (3.32 ± 0.27) compared to PBS controls (1.69 ± 0.35) (p<0.05, n=4) (Figure S2E). This change was not seen in cells of lymph nodes of these mice (Figure S2F).

**Figure 3.**
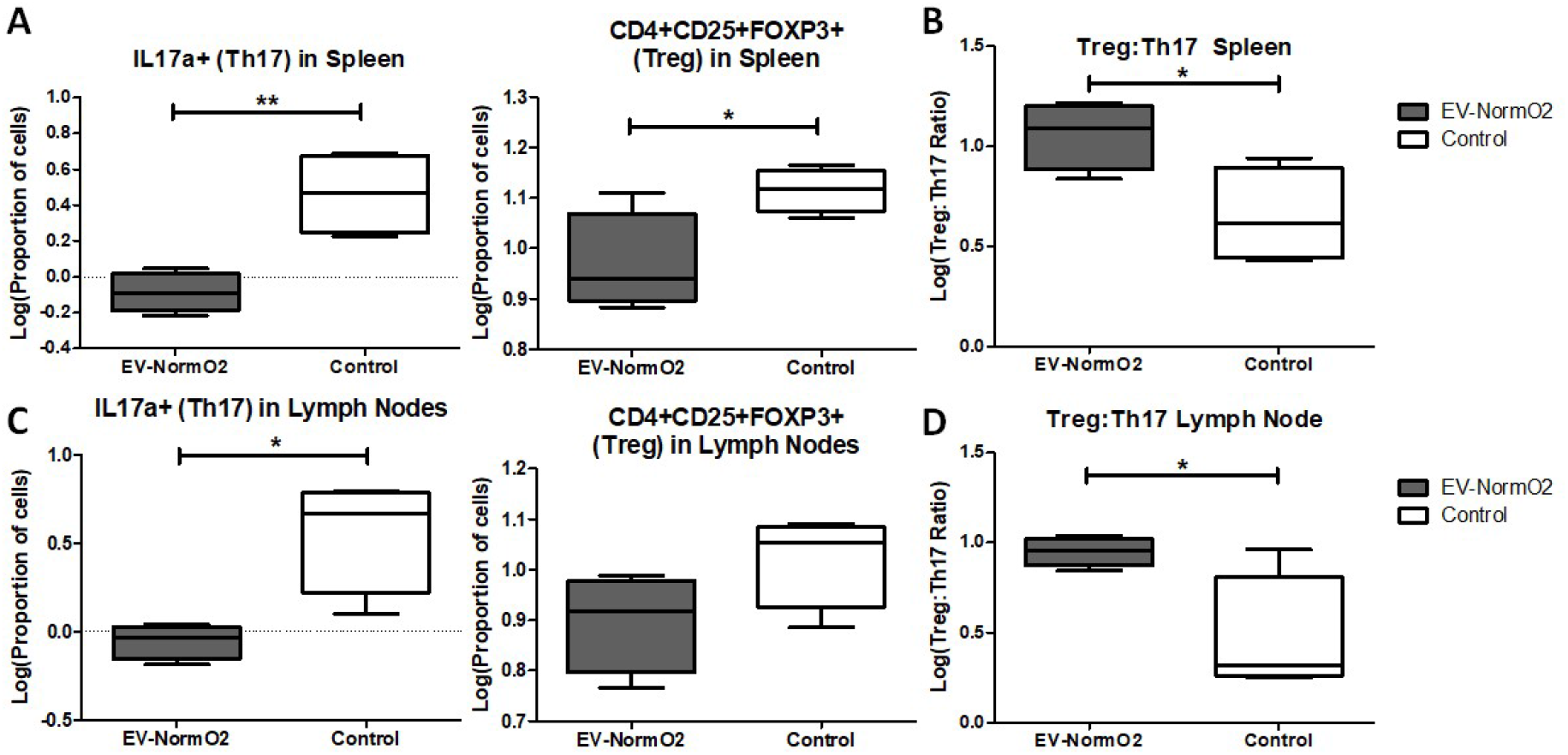
Outcomes of intracellular staining for T cell polarisation analysis. Intracellular staining for FACS analysis of IFN-γ, IL4 and IL17a in CD4+ T cells from EV-NormO2 versus PBS control. Significant reductions in (A/C) IL17a expression suggestive of reduced Th17 polarisation in Spleen and Lymph Node with a small or insignificant change in regulatory T cell polarisation (B/D) resulted in trend for restoration of the Treg:Th17 balance in EV-NormO2 treated mice which was found significant in lymph node cells only (n=4, *p<0.05; **p<0.01)(Unpaired T Test using log transformed data).

The outcomes of treatment using EVs *in vivo* demonstrate a reduction in polarisation towards IL17a secreting T helper cells for both spleen and lymph nodes of AIA treated mice that is not seen *in vitro* using EV-treated activated T cells from healthy mice (without AIA induction). We therefore hypothesise that the therapeutic effect seen *in vitro* may require the presence of a mediator and we propose the candidate for this would be DCs. Regulatory T cells influence DCs through EVs mediated intercellular communication to provide immunomodulation, including regulation of tolerance in autoimmune conditions^24^. Potentially the variation observed through *in vivo* treatments compared with *in vitro* responses reflect the absence of mediating cell types such as DCs.

## Discussion

This study builds on the growing knowledge for the therapeutic efficacy of EVs as a treatment for inflammatory autoimmune disorders. Here, we demonstrated the first application of MSC-derived EVs into the AIA model of inflammatory arthritis with examination of both hypoxic and pro-inflammatory cell priming evaluated against standard EV generation. For this investigation, the AIA model of inflammatory arthritis is pertinent in investigating the T cell mediated inflammatory response, as AIA is specifically driven through CD4+ T-lymphocyte responses leading to synovial leukocyte infiltration^35^, in comparison to the more commonly applied collagen-induced arthritis (CIA) model which involves a breach of immune tolerance and generation of systemic polyarticular disease through production of autoantibodies leading to synovitis^36^. Consequently, whilst CIA has recently been applied to evaluate the efficacy of EVs therapies as a model of RA, producing results demonstrating strong immunomodulatory effects with unknown mechanism^19^ or indicating T-lymphcyte mechanisms underpin therapeutic outcomes^37^, the AIA model remains appropriate for examining molecular changes evoked through the immunomodulatory action of EVs and their impact upon CD4+ T cells^36^ as shown in this study. We aim to inform researchers and clinicians on the efficacy of priming strategies to prepare EVs as an enhanced therapy. In our study, EV treatments prompted amelioration of clinical symptoms of AIA with reduction of joint swelling both with and without priming. Examination of joint sections showed improved histological features following standard and pro-inflammatory primed MSC-derived EV treatments compared to controls but not when using hypoxically derived EVs. Mechanistically, EVs acted remarkably different from their cells of origin (MSCs) when co-cultured with CD4+ T cells from healthy murine spleens. Standard and hypoxically-derived EVs reduced Th17 polarization without affecting T cell proliferation, while MSCs increased Th17 polarization, and lessened T cell proliferation. Our previous study indicated that MSCs do indeed increase Th17 polarisation but this is offset by increases in Treg cells to restore a more homeostatic balance between the two. Here we examined the Treg:Th17 ratio following AIA initiation and treatment with EV-NormO2. With lack of significant improvements in hypoxic priming outcomes, we opted to test only EV-NormO2 for influence on T cell polarisation following treatment application in the pro-inflammatory environment. *Ex vivo*, CD4+ T cells isolated from spleens of arthritic mice treated with standard EV-NormO2 again showed significantly reduced Th17 polarization that rebalanced the Treg:Th17 ratio. Together, this suggests the reduction in Th17 cells led to restoration of the Treg:Th17 ratio, which is typically unbalanced in inflammatory arthritis, and reduced immune cell recruitment. This result was accompanied with an increase in IFN-γ expressing T cells (Th1) and concomitant increase in the Th1:Th2 ratio which should be taken into account when considering clinical interventions, although for this study the interest was directed towards RA therapies targeting Treg:Th17 ratios.

To further dissect the therapeutic mechanism of action of MSC-derived EVs, circulating levels of TNF-α, which is a key driver of pathogenesis in RA and therapeutic target in biological treatments, and IL-10, which is a master regulator of anti-inflammatory immune responses, were measured in mice with AIA at day 3. While IL-10 was not detected in our assay, similarly low levels of TNF-α were detected in serum of untreated and EV-treated mice. However, whilst this suggests that TNF-α blockade and IL-10 modulation are not the mechanisms by which MSC-derived EVs improve AIA, it is noted that circulating serum levels of TNF-α rise rapidly in 24 hours post-induction then fall rapidly in the AIA model, so detectable serum TNF-α may not be significantly different in AIA and control mice at 3 days post-induction^38,39^. This mechanism may be more suited to assessment following prophylactic treatment using EVs administered at the point of induction rather than after onset of symptoms.

Our previous study demonstrated reductions in Th17 cells following CM-MSC treatment compared to control and MSC treatments. Whilst EVs have been shown to increase Il-10 in macrophages^40^, this study indicates that the presence of MSCs, and not just their secretome, may be driving increased IL-10 expression influencing T cell polarisation *in vivo*, and were responsible for the reduction in Th1 and increase in Th2 cells seen in our previous study^3^. In this study we observe an increase in Th1-like cells following EVs treatments. EVs are capable of significantly reducing Th17 cell numbers, however, and this represents a significant finding in the search for immunomodulatory therapeutics for treating autoimmune disorders where an imbalance in T cell polarisation (Treg:Th17) is integral in disease pathology. Moreover, EV treatment in this study resulted in an improved (2.40-fold over controls) Treg:Th17 ratio compared to our previous results using whole CM-MSC treatment (2.13-fold over control) or MSC treatment (1.47-fold over control)^3^.

The proportions of Treg and Th17 cells in RA sufferers has been directly linked to the severity of disease, and restoring the Treg:Th17 balance has the potential to promote homeostasis and positive clinical outcomes. Here, we demonstrate that untreated (PBS control) spleens of mice with AIA display a Treg:Th17 ratio of 5:1 and that this is improved to 12:1 upon EV-NormO2 treatment. This result reinforces the use of vesicular secretome as a therapeutic option for RA treatment. Furthermore, MSC-EVs have been shown previously to be the component of the secretome responsible for *in vitro* immunosuppression of activated T cell responses^41,42^ and our study both replicates and builds on these results through extrapolation to the *in vivo* environment and specific examination of T cells from spleen and lymph nodes isolated from *in vivo* experimentation. We additionally offer data to support the selection of standard culture practices as advantageous in producing broad immunomodulatory outcomes, in comparison to either hypoxic or pro-inflammatory priming of MSCs for EVs production.

We observed effects on T cell polarisation that were consistent between *in vitro* and *ex vivo* outcomes, with the exception of reduced Th2 cells *in vitro* being unmatched *ex vivo.*

It has been suggested that the immunomodulatory properties of EVs are contingent on pro-inflammatory priming^19,43^. However, the present work demonstrates that whilst priming strategies may enhance some therapeutic outcomes, the mechanisms are unclear and the overall immunomodulatory property may be inherent to MSC-EVs regardless of priming methodology. This would suggest the benefits of priming strategies for EVs production may be dependent on MSC variability. EV-Pro-Inflam were the least effective treatment methodology for T cell deactivation, however they did reduce swelling and immune cell recruitment to the synovium. We hypothesise that these effects are due to alterations in the EV cargo prompted by the culture conditions, with our previous studies demonstrating modulations such as increased chemotaxis and angiogenesis prompted through pro-inflammatory priming of MSCs prior to EVs collection^44^. In contrast, hypoxically primed MSCs produced EVs that reduced swelling but failed to have any impact upon the histological outcomes measured in AIA joints. Neither priming strategy demonstrated consistent improvements in immunomodulatory or tissue regenerative outcomes.

Our results suggest that the *in vivo* response to pro-inflammatory primed EV treatment in the AIA model of inflammatory arthritis is not primarily through inhibition of T cell activation, but through suppression of CD4+ Th17 effector polarisation combined with reduction in synovial infiltration in the affected joint. Although this effect is more pronounced following pro-inflammatory priming of MSCs in comparison to standard EV production, the reduction in exudate and hyperplasia seen with EV-NormO2 treatments is likely to convey a greater therapeutic response. Pannus tissue is thought to develop from cells of the synovial membrane and cells infiltrating the joint cavity in response to pro-inflammatory signalling^45,46^. The reduction of synovial infiltrate, hyperplasia and synovial exudate observed in EV-NormO2 treatments *in vivo* may therefore translate to reduced pannus formation. These benefits, in combination with improved Th17:Treg balance would convey advantageous outcomes in comparison to either priming methodology.

We hypothesise that whilst the overall effectiveness of these treatment methodologies is similar in potency, the outcomes differ due to variations in underlying mechanisms of action. Our results demonstrate hypoxic preconditioning reduces swelling *in vivo* and affects Th17 polarisation of T cells *in vitro*, however insufficient evidence was found to propose a clear mechanism for therapeutic capacity. MSCs under hypoxic culture have been shown to promote anti-inflammatory M2 macrophage polarization^47^, reduce reactive oxygen species (ROS) and upregulate TGF-β, IL-8, IL-10 and PGE2, which are also implicated in macrophage polarisation and MSC immunomodulatory capacity^47–49^. Consequently, we hypothesise that effects seen *in vivo* following hypoxic preconditioning may be related to reduction in macrophage mediated immune responses and more appropriate for tissue regenerative applications. Furthermore, hypoxic culture of MSCs has been shown to promote vascularisation^50,51^ which, whilst beneficial for tissue repair, may actually contribute to pannus formation in the RA joint^46^. Given these considerations, the evidence here suggests that normal culture conditions are sufficient to elicit an optimal response in MSCs for EVs production ahead of therapeutic application into the inflammatory environment.

Previously, we determined that joint swelling in induced arthritis as a clinical indication of joint inflammation is reduced in the presence of administered MSCs and these cells migrate into the inflamed synovium^2^. Joint swelling is common with different types of arthritis and is caused by oedema due to the endothelial cells of blood vessels becoming leaky in the inflamed synovium. We hypothesised that soluble factors produced by MSCs are responsible for permeability changes in the synovial endothelial cells as these cells did not co-localise, although further studies are required in this regard. The therapeutic potential of MSC-derived EVs has been documented in an animal model of radiation-induced lung injury where EVs attenuated radiation-induced lung vascular damage, inflammation, and fibrosis via miRNA-214-3p^52^ and in a rodent model of haemorrhagic shock-induced lung injury model where EVs attenuated pulmonary vascular permeability and lung injury^53^.

Given the observed outcomes, we hypothesise that the similarities in immunomodulation between EV-2%O2 and EV-Pro-Inflam treatment may lie in their influence on STAT3 activation. Increased HIF-1α activates STAT3^54^. STAT3 induces RORγt expression and is activated by TNF-α, IL-6, IL-21 and IL-23, which are key cytokines in promoting Th17 differentiation, and RORγt is a master regulator of Th17 differentiation^55,56^. Moreover, STAT3 activation has been shown to be necessary for the development of Th17 cells and Th17 related autoimmunity, such as seen in RA^56^ so EVs inhibiting STAT3 activation could lead to the drop in Th17 polarisation observed. EVs from other mesenchymal cell types are implicated in influencing STAT3 signalling to induce mesenchymal transition^57^ and EVs from macrophages are able to trigger nuclear translocation of STAT-3^40^, demonstrating STAT3 activation as a target of EVs influence. STAT3 has been suggested as a therapeutic target in the treatment of autoimmune inflammatory disorders such as RA^58^ and inhibition of STAT3 by EVs would have direct therapeutic potential. Furthermore, STAT3 is implicated in interferon responses, which may present a potential mechanism contributing to the increased IFN-γ expressing T cells observed following EV-NormO2 treatments *in vivo.* Further work can elucidate this mechanism of action.

In this study, our primary aim was to evaluate the efficacy of EV treatments *in vivo* during AIA. We show that MSC priming leads to the release of EVs that if administered to mice with acute inflammatory arthritis significantly ameliorate disease pathogenesis, mainly through inhibition of Th17 polarization. Future studies will define the composition and sub-vesicular localisation of proteins in EV cargos.

### Conclusions

This study provides new evidence that supports the use of EVs in clinical therapies for RA and similar autoimmune disorders. The possibility to manipulate the protein and nucleic acid cargo of EVs through control of parental MSC cultures offers a novel opportunity for targeted therapies that can be tailored to individual pathological features of RA, advancing personalised medicine. Further work on the control of EV cargo will elucidate the molecular mechanism of action underlying varied responses between priming strategies and assist in the efficacy of cell-based therapies in the clinic. This study aims to support the growing body of evidence for the introduction of EVs into the therapeutic milieu.

## Methods

### Cells and EVs

Primary human MSCs were isolated from commercially available bone marrow aspirate (Lonza, USA) using an adherence technique^48^ and each donor material cultured in both normoxic (21%O2) or hypoxic (2%O2) conditions from isolation of MSCs in Dulbecco’s Modified Eagles Medium (DMEM) with 10% foetal bovine serum (FBS) and 1% penicillin-streptomycin (n=3 aspirates). Hypoxic cell culture was achieved using 2% O2 in hypoxic workstation (InvivO2 Physiological Cell Culture Workstation, Baker Ruskinn) to ensure cells were not exposed to environmental oxygen levels at any stage of isolation or culture. Cells (P3-P5) were characterised as MSC through immunophenotyping of surface markers with flow cytometry and tri-lineage differentiation^48^. Isolations of EVs were performed over a 48 hour culture period from MSCs in normoxic 21% oxygen culture (EV-NormO2, n=11 isolations); MSCs in hypoxic 2% oxygen culture (EV-2%O2, n=4 isolations) and from MSCs previously cultured for 48 hours with pro-inflammatory cytokine cocktail comprising interferon-gamma (IFN-γ, 10ng/ml), tumour necrosis factor-alpha (TNFα, 10ng/ml) and interleukin 1 beta (IL-1β, 5ng/ml) prior to transfer to serum-free medium for 48 hours for EV collection (EV-Pro-Inflam, n=4 isolations).

### Differential ultracentrifugation for isolation of EVs

MSCs from the same batch used for cell treatment (P3-P5) were cultured to 80-90% confluence in T75 flasks, washed with PBS three times and then serum-free DMEM. Flasks were incubated for 48 hours with 12ml serum-free DMEM at 37°C, 5% CO2. After 48 hours, CM-MSC was removed and a cell count performed to determine the number of cells used to produce EVs. EVs were isolated from CM-MSC by differential ultracentrifugation. In brief, CM-MSC was centrifuged for 10 minutes at 300 *x g* to remove cell debris then again for 10 minutes at 2000 *x g* to remove apoptotic bodies and residual dead cells. Supernatant was again taken and centrifuged at 10,000 *x g* for 45 minutes in an ultracentrifuge (Beckman Coulter) to remove larger/denser EVs. Supernatant was again retrieved and passed through a 0.22μM syringe filter. Filtered supernatant was then centrifuged using fixed angle type 70.1Ti rotor (Beckman Coulter) in a Beckman L8-55M ultracentrifuge, k-factor 122.6 at 100,000 *x g* for 90 minutes to isolate a pellet comprising small EVs. The EVs pellet was resuspended in 5ml of PBS to wash EVs and then spun again at 100,000 *x g* for 60 minutes, the supernatant discarded and the residual EV pellet resuspended in PBS at 30μL per 1.0 × 10^6^ cells used in EV generation and stored at 4°C to be used within 24 hours or at −20°C for later use.

### Characterisation of EVs

Successful isolation of EVs was confirmed through multiple methods in accordance with ISEV guidelines^59^. Samples of vesicle preparations underwent BCA for total protein concentration in EV preparations following manufacturer’s instructions with EV disruption in a sonicating water bath (30 seconds sonication at 1 minute intervals for 3 cycles) to release intracellular protein and assess cargo proteins as well as surface proteins. Data passed a Kolmogorov-Smirnov test for normality and was analysed with a repeated measures 1 way ANOVA with Tukey’s post hoc multiple comparison test.

Vesicles were examined using flow cytometry for expression of characteristic markers CD9, CD63 and CD81 (Miltenyi MACSPlex Exosome Identification Kit, human) following manufacturer’s instructions for analysis. Particle by particle analysis of vesicle size was assessed using Nanopore technology (Izon Science Ltd) tuned in the region ~80-300nm, to confirm the size fractions being applied experimentally.

Images of EVs were obtained using TEM. EVs to be imaged by TEM were isolated as detailed above, however residual EV pellet after PBS wash was resuspended in 30μl Milli-Q water per 3.0 × 10^6^ cells. The resulting EV suspension was spotted onto a glow discharged holey carbon mesh copper grids (Quantifoil, R2/1) and incubated at room temperature for 4 mins 45 seconds before 5μL of 0.22μm filtered 2% (v/v) uranyl acetate was added. This was left to incubate at room temperature for a further 90 seconds before the excess liquid was removed using blotting paper. Grids were then imaged using a FEI Tecnai G2 12 Biotwin (LaB6, accelerating voltage 100kV).

EV protein content was examined by Western Blotting to identify the presence of Alix and absence of mitochondrial Cytochrome C (CYC1) in isolated particles, as per the minimal criteria for identification of EVs^60^. Briefly, ten micrograms of EVs and cells were lysed with 2x Laemlli buffer with β-mercaptoethanol and denatured by heating at 70°C for 5min. Samples were electrophoresed on a 4-12% TGX stain free gel (Bio-Rad, CA, USA) at 200V for 30-40min. Samples were then blotted onto a nitrocellulose membrane using the wet transfer method overnight at 100V. After blotting, membranes were blocked for 2 hours in 5% semi-skimmed milk in Tris-buffered saline with 0.05% Tween (TBST) followed by an incubation in primary antibodies Alix (1:200), CYC1 (1:200) (Santa Cruz Biotechnology, TX, US) for 2 hours at RT. After three five-minute washes, the membrane was probed with a goat anti-mouse IgG-HRP conjugated secondary antibody (1:1,000 in TBST, Life Technologies Limited, Paisley, UK). HRP-conjugated secondary antibody was added for 1h and the wash step repeated. SuperSignal West Femto Chemiluminescent Substrate (Thermo Fisher Scientific, Waltham, MA, USA) was added to the membrane and imaged with ChemiDoc™ Touch Imaging System (Bio-Rad, CA, USA) using Image Lab v.6.0.1 software (Bio-Rad, CA, USA).

### Co-culture of healthy T cells with EVs

Proliferation of activated T cells was assessed as a measure of T cell deactivation. Initially, for positive controls 5×10^4^ MSCs per well were cultured in 96-well plates for 24 hours at 37°C. CD4+ T cells were purified from spleens and lymph nodes (popliteal/inguinal) of healthy C57Bl/6 mice using the CD4+ T Cell Isolation Kit (Miltenyi) following manufacturer’s instructions. T cells were seeded at a density of 5.0×10^5^/well in 250μl RPMI medium with 10% FBS and cultured for 5 days with MSCs (ratio 10:1) or with EV-NormO2, EV-2%O2 or EV-Pro-Inflam (ratio equivalent to secretions from 10:1 cells) (n=8). T cells alone served as control. Cells were activated using anti-Biotin MACSiBead Particles (Miltenyi) (ratio 2:1). EV treatments were refreshed after 2 days of culture. Polarisation was assessed as above and proliferation measured through reduction in signal intensity using VPD450 Violet proliferation dye (BD Biosciences)^3^. Data was analysed using a 2 way repeated measures ANOVA with Bonferroni multiple comparison test post hoc.

### Antigen-induced arthritis (AIA) model of inflammatory arthritis

Animal procedures were undertaken in accordance with Home Office project licence PPL40/3594. AIA was induced in male C57Bl/6 mice (7-8 weeks) as previously described^2,3,61^. Swelling was assessed by measuring the difference in diameter between the arthritic (right) and non-arthritic (left) knee joints (in mm) using a digital micrometer (Kroeplin GmbH) before and at set time points after treatment. Independent experiments were performed to assess swelling and histopathological effects of EV-NormO2 (n=10), EV-2%O2 (n=6) and EV-Pro-Inflam (n=6) compared with PBS controls (n=21) with all EVs sourced from matching MSCs and statistical analysis performed using 2 Way ANOVA for Repeated Measures with Bonferroni Multiple Comparisons Test post hoc; or for *in vivo* primed T cell collections using EV-NormO2 compared with PBS control (n=4) and log transformed data analysed with Unpaired T tests. Peak swelling was observed at 24 hours post-induction and results at subsequent timepoints are expressed in millimetres as reduction from peak swelling.

### Intra-articular injection of EVs

Treatments comprising 15μl of EVs suspension in PBS corresponding to EV secretions from ~5.0 × 10^5^ cells or PBS alone controls were injected intra-articularly 1 day post arthritis induction with 0.5ml monoject (29G) insulin syringes (BD Micro-Fine, Franklyn Lakes, USA) through the patellar ligament into the right knee joint. Joint diameters were measured at 1, 2 and 3 days post injection. Blood, joints, spleen, inguinal and popliteal lymph nodes were collected immediately post-mortem. Four independent experiments were performed to assess primed EVs impact on joint swelling and histopathology (Control n=21, EV-NormO2 n=10, EV-2%O2 and EV-Pro-Inflam n=6). A further independent experiment was conducted to address the potential for EV immunomodulation of T cells in the *in vivo* environment in AIA (EV-NormO2 versus PBS controls, n=4 per condition). All measures were taken to reduce animal numbers wherever possible.

### Arthritis Index

Animals were sacrificed for histological analysis at day 3 post arthritis induction. Joints were fixed in 10% neutral buffered formal saline and decalcified in formic acid for 4 days at 4°C before paraffin embedding. Sections (5μm) were stained with haematoxylin and eosin (H&E) and mounted in Hydromount (National Diagnostics) as described previously^2,3^. H&E sections were scored for hyperplasia of the synovial intima (0=normal to 3=severe), cellular exudate (0=normal to 3=severe) and synovial infiltrate (0=normal to 5=severe) by two independent observers blinded to experimental groups^62^. Scores were summated, producing a mean arthritis index. Data was analysed using a 1 way repeated measures analysis of variance test with Bonferroni multiple comparison test post hoc.

### Cytokine Quantification

IL-10 and TNFα in serum and CM-MSC were quantified using mouse Quantikine ELISA IL-10 immunoassay (R&D Systems) and TNFα ELISA high sensitivity (eBioscience) respectively, following manufacturer’s instructions.

### T cell polarisation

Spleens and lymph nodes (popliteal/ inguinal) were collected from mice 3 days post-arthritis induction and dissociated as described previously^3^. Splenocytes and pooled lymph node cells were seeded separately at 1.0×10^6^ cells/well in 96-well plates (Sarstedt) in RPMI-1640 with 10% FBS, 0.05μg/mL IL-2 and activated with cell stimulation cocktail (eBioscience) for 1 hour prior to adding 10μg/ml brefeldin A (Sigma) and culturing for a further 4 hours. Unstimulated T cells and cells without brefeldin A served as negative controls. Following activation, cells were resuspended in 2mM EDTA in PBS and Tregs (CD4+CD25+FOXP3+) were isolated using the CD4+CD25+ Regulatory T Cell Isolation Kit (Miltenyi) following manufacturer’s instructions; or stained for T cell subset identification. For this, cells were permeabilised using permeabilisation buffer kit (eBioscience) and intracellularly stained with anti-mouse IFN-γ (Th1), IL-4 (Th2) or IL17a (Th17) (eBioscience). Cells were analysed on a BD FACS Canto II flow cytometer and comparisons drawn for percentage CD4+ cells and signal intensity (XGeoMean) for each antibody.

### Statistical analysis

Data were tested for equal variance and normality using D’Agostino & Pearson omnibus normality test. Differences between groups were compared using 1-way ANOVA with Bonferroni post hoc for parametric data or Kruskal-Wallis ANOVA with Dunn’s post hoc test for non-parametric, or 2-way ANOVA with Bonferroni correction, as stated. Repeated measures tests using log transformed data was applied to Protein concentration and analysed with log transformed data using a 4 parameter polynomial nonlinear regression to interpolate results from a standard curve. *In vivo* comparisons of EV-NormO2 treatments versus PBS controls were analysed by log transformation of percentage data and comparison using unpaired T tests. All statistical analysis was carried out using Prism 5 (GraphPad software) or IBM SPSS Statistics 24.0, with P<0.05 deemed statistically significant. Results are expressed as mean ± standard error of the mean using *p<0.05, **p<0.01, ***p<0.001.

## List of abbreviations

AIA: Antigen Induced Arthritis model
BCA: Bicinchoninic Acid Protein Assay
CM-MSC: MSCs derived conditioned medium
CTSD: Cathepsin D
CYC1: Cytochrome C
DMEM: Dulbecco’s Modified Eagle’s Medium
EV-2%O2: EVs derived from cells cultured in 2% oxygen
EV-NormO2: EVs derived from cells cultured in normal environmental oxygen ~21%
EV-Pro-Inflam: EVs derived from MSCs cultured with pro-inflammatory cytokine stimulation
EVs: Extracellular vesicles
FBS: Foetal bovine serum
FIH: Factor inhibiting Hif-1 protein
H&E: Haematoxylin and eosin stain
HIF-1α: Hypoxia inducible factor 1 alpha
ICAM-1: Intercellular adhesion molecule 1
IDO: Indoleamine 2,3-dioxygenase
IFN-γ: Interferon gamma
IL-10: Interleukin 10
IL-1β: Interleukin 1 beta
ISEV: International Society of Extracellular Vesicles
LFA-1: Leukocyte function associated antigen 1
MFI: Mean fluorescence intensity
miRNA: micro ribonucleic acid
MMPs: Matrix metalloproteins
MSCs: Mesenchymal stem/stromal cells
MVBs/MVEs: Multivesicular bodies/endosomes
NO: Nitric oxide
PBS: Phosphate buffered saline
RA: Rheumatoid Arthritis
RF: Rheumatoid factor
RISC: RNA-induced silencing complexes
RORγt: Retinoic acid receptor-related orphan receptor-γt
ROS: Reactive oxygen species
TIMPs: Tissue inhibitor of metalloproteinases
TNF-α: Tissue necrosis factor alpha
Treg: Regulatory T cell
VEGF: Vascular endothelial growth factor
VHL: von Hippel-Lindau ubiquitin ligase

## Acknowledgements

The authors would like to thank the Nanoscale and Microscale Research Centre, University of Nottingham, for access to TEM facilities. We would like to thank Dr. Mark Platt (Loughborough University, UK) for his assistance in Nanopore technology.

## Author’s contributions

A.K. performed experimental work, wrote the manuscript and prepared all figures. K.T. provided support and technical skills for T cell investigations. P.R. gave technical expertise on Nanopore technology. R.M. provided support for histological investigations, TEM experiments and discussed and commented on the manuscript. R.L. performed TEM experiments. M.H. performed ELISAs and provided support for *in vivo* work and histological experiments. A.M.P. gave technical advice on data interpretations and presentation and reviewed the manuscript. N.R.F. contributed to experimental design. O.K. conceived and designed the research study, performed *in vivo* experiments, analysed data, wrote and supervised the manuscript. All authors reviewed the manuscript before submission.

