## Supplementary Figures S1 and S2 for "THERAPEUTIC EFFECTS OF HYPOXIC AND PRO-INFLAMMATORY PRIMING OF MESENCHYMAL STEM CELL-DERIVED EXTRACELLULAR VESICLES IN INFLAMMATORY ARTHRITIS"

**Kay et al 2021 Supplementary Figures**

Figure S1

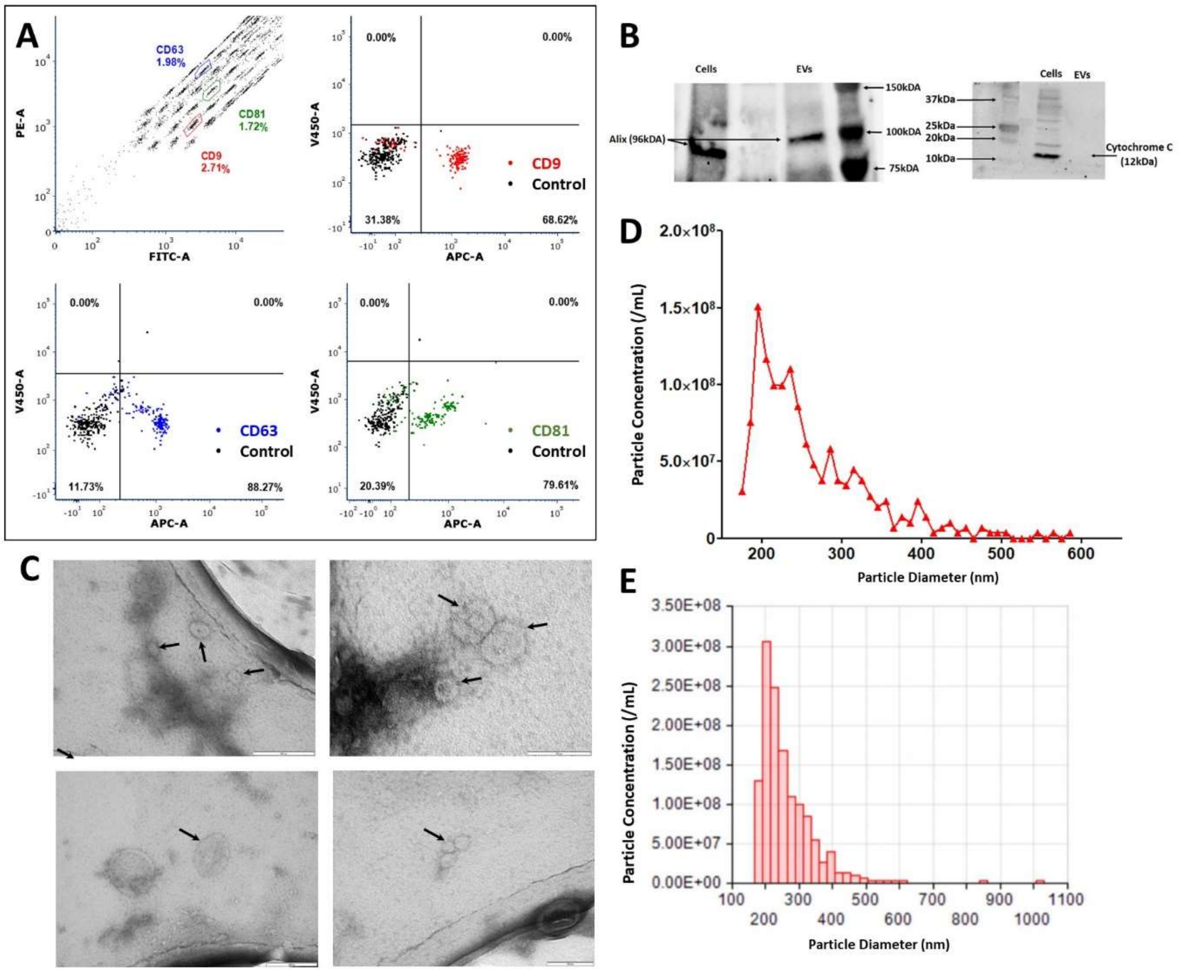


**Figure 1: Detailed characterisation of EVs.** (A) Representative flow cytometry analysis of EV-NormO2 preparations using MACSPlex exosome detection kit (Miltenyi) for the detection of CD9 (mean 81.25±5.03); CD63 (mean 94.59±2.23); and CD81 (mean 79.41±9.07) with unstained control beads. (B) Western blotting demonstrates presence of Alix and absence of cytochrome C in EVs (C) TEM characterisation of hBM-MSC derived small EVs. Small EVs were re-suspended in sterile distilled water after isolation and spotted onto TEM grids before being stained with uranyl acetate. Black arrows indicate recorded small EVs. Small EVs were isolated from conditioned media taken from hBM-MSCs isolated from bone marrow aspirate cultured in hypoxic conditions (EV-2%O2). (D) (E) Representative output from particle concentration and EV sizing Nanopore analysis (Izon) tuned in the region ~80-300nm, highlighting EVs diameter range, with peak diameter averaging around 200nm with maximal diameter around 500nm (n=11).

Figure S2

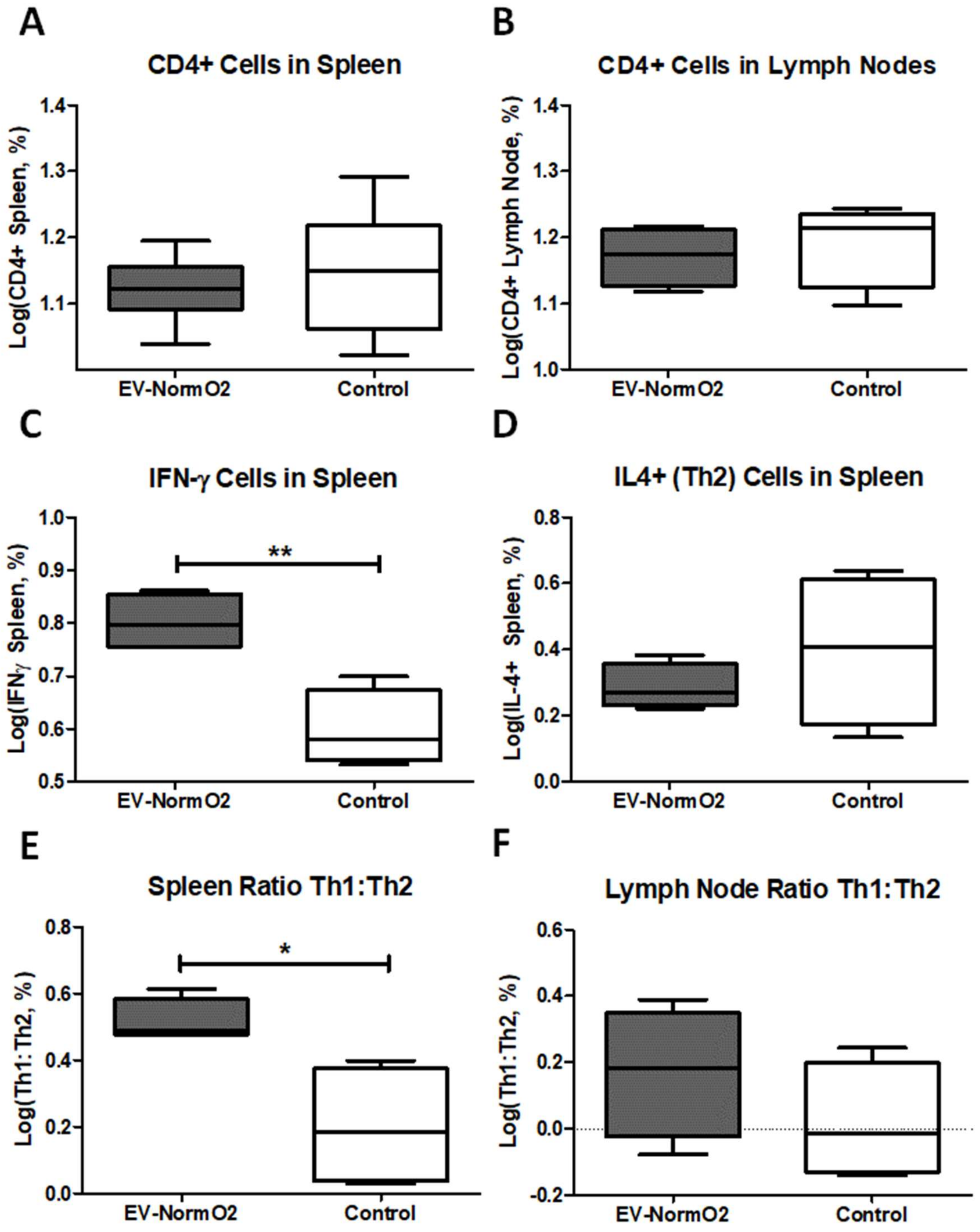


**Figure S2 – Outcomes of intracellular staining for T cell polarisation analysis.** Intracellular staining for FACS analysis of IFN-γ, IL4 and IL17a in CD4+ T cells from EV-NormO2 or PBS control spleen and lymph node tissues. (A) EV-NormO2 treatments showed no change in CD4+ cell presence in spleen or lymph node tissue compared to PBS controls. (C/D) A significant increase in IFN-expressing (Th1) cells was observed in spleens with no change in IL-4 expressing (Th2) cells (E/F) leading to an increase in the Th1:Th2 ratio in spleen, but not lymph node, tissues (n=4, *p<0.05; **p<0.01)(Unpaired T Test using log-transformed data).
